# In vivo imaging and quantification of carbon tracer dynamics in nodulated root systems of pea plants

**DOI:** 10.1101/2021.06.23.449643

**Authors:** Ralf Metzner, Antonia Chlubek, Jonas Bühler, Daniel Pflugfelder, Ulrich Schurr, Gregor Huber, Robert Koller, Siegfried Jahnke

**Affiliations:** IBG-2: Plant Sciences, Forschungszentrum Jülich GmbH, Jülich, Germany; Biodiversity, University of Duisburg-Essen, Universitätsstr. 5, 45141 Essen, Germany

**Keywords:** ^11^C, Biological Nitrogen Fixation, Legumes, Nitrate inhibition, PET, Radiotracer

## Abstract

Legumes associate with root colonizing rhizobia that provide fixed nitrogen to its plant host in exchange for recently fixed carbon. There is a lack in understanding how individual plants modulate carbon allocation to a nodulated root system as a dynamic response to abiotic stimuli. One reason is that most approaches are based on destructive sampling, making quantification of localized carbon allocation dynamics in the root system difficult. We established an experimental workflow for routinely using non-invasive Positron Emission Tomography (PET) to follow the allocation of leaf-supplied ^11^C tracer towards individual nodules in a three-dimensional (3D) root system of pea (*Pisum sativum*). Nitrate was used for triggering the shutdown of biological nitrogen fixation (BNF) expected to rapidly affect carbon allocation dynamics in the root-nodule system. This nitrate treatment lead to a reduction of ^11^C tracer allocation to nodules by 40% – 47% in 5 treated plants while the variation in control plants was less than 11%. The established experimental pipeline enabled for the first time that several plants could consistently be labelled and measured using ^11^C tracer in a PET approach to quantify C-allocation to individual nodules following a BNF shutdown. This demonstrates the strength of using ^11^C tracers in a PET approach for non-invasive quantification of dynamic carbon allocation in several growing plants over several days. A major advantage of the approach is the possibility to investigate carbon dynamics in small regions of interest in a 3D system such as nodules in comparison to whole plant development.

**One sentence summary:** Positron Emission Tomography for quantification of carbon allocation dynamics in individual nodules within a 3D root system revealed strong effect of nitrate on carbon allocation.

## Introduction

Legumes establish a symbiosis with nitrogen (N)-fixing rhizobia. In specialized root nodules, rhizobia are supplied with plant derived carbon (C) and in return, rhizobia provide the plant with fixed, atmospheric N (Gordon et al., 1999, Graham and Vance, 2003). Legumes play a key role in natural ecosystems and agriculture, as they do not require additional application of synthetic N fertilizers and produce grains with high protein content for humans and cattle (Rubiales and Mikic, 2015, Stagnari et al., 2017). Nodules depend to a large part on recently fixed carbon from the leaves (Kouchi et al., 1986, Fujikake et al., 2003). A plant host may invest up to 30% of plant photosynthates into his symbiont (Provorov and Tikhonovich, 2003). Consequently, a timely and balanced carbon allocation in return for N is vital for plant performance. While much progress has been made associated with regulation of biological N fixation (BNF) at the level of transporters (Valkov et al., 2017) and gene expression (Cabeza et al., 2014), the overall mechanism and the linkage of external N-fertilization to dynamic carbon allocation remains unclear (Schulze, 2004, Schwember et al., 2019).

One reason for the lack of understanding in host-rhizobia interactions is the difficulty of studying carbon allocation in vivo. The root tissues involved are especially sensitive to manipulation, not easily accessible belowground, and nodules are formed in the complex 3D structure of the root system. Carbon tracer approaches are well suited for carbon allocation studies because tracers can be administered non-invasively as CO_2_ to single leaves or the whole shoot. Carbon tracers have been successfully employed for disentangling carbon allocation between plants and nodules mostly using the stable isotope ^13^C (Voisin et al., 2003) or the long-lived radioisotope ^14^C (Small and Leonard, 1969, Vessey et al., 1988). Both isotopic approaches require destructive harvesting of the plant material, preventing studies on the dynamics of carbon in a nodulated root system of individual plants. In contrast, the short-lived carbon radioisotope ^11^C (t_1/2_ =20.4 min) allows for carbon tracer studies in vivo and enables detailed time series of ^11^C tracer allocation processes at multiple locations along the shoot-root axis and according to dynamic abiotic and biotic conditions (Thorpe et al., 1998, Minchin and Thorpe, 2003, Henkes et al., 2008, Mason et al., 2014). However, previous approaches to monitor carbon partitioning in nodulated root systems using ^11^C were based on individual scintiallation detectors (Thorpe et al., 1998, Walsh et al., 1998a, Walsh et al., 1998b) covering extended sections of plants integrating e.g. all roots and did not permit to analyse the complex 3D structure of a nodulated root system in detail. Further, most of the approaches were very limited in numbers of biological replicates due to delicate experimental set-ups. The application of ^11^C in combination with Positron emission tomography (PET) has the potential to overcome these limitations. PET allows for a non-invasive 3D imaging of whole plants and enables to visualize the transport and distribution of ^11^C tracer within a plant system (Jahnke et al., 2009, De Schepper et al., 2013, Hubeau and Steppe, 2015).

While the physics and technical aspects of the PET instruments are more complex than for scintillation detectors set-ups, PET scanners are routinely applied as a medical diagnostic tool. It has been demonstrated that medical PET scanners could even directly be employed for plant studies (Karve et al., 2015, Ruwanpathirana et al., 2021). A pilot study with a 2-dimensional PET sytem showed the principal feasability of detecting ^11^C tracer in nodules (Fujikake et al., 2003), but did not allow for the analysis of local carbon allocation to individual nodules or within a longer time period. The combination of PET using ^11^C labelling and Magnetic Resonance Imaging (MRI) resulted in a high spatial and temporal resolution of ^11^C tracer transport processes and showed the potential for studying dynamic carbon allocation to a local region of interest within a complex 3D root system (Jahnke et al., 2009). However, for using the full potential of PET in plant sciences a framework for routine quantification of ^11^C tracer allocation is still missing.

Here, we report the set-up of such a concept of repeated PET measurement of individual plants for quantifying ^11^C tracer dynamics in nodulated root systems of pea plants. Plants were labeled with ^11^CO_2_ twice a day over the course of four days. A nitrogen (nitrate) treatment was applied in the root zone before the third day of measurements. The guiding question was if the PET set-up enables detection of localized responses to abiotic stimuli, i.e., whether changes in ^11^C allocation to the nodules can be observed after administering nitrate to the roots?

## Results

### The experimental Pipeline: Overview

We established a measurement workflow for repeatable non-invasive measurements of short-term carbon allocation with ^11^C-PET in nodulated root systems **(Fig. 1**). It consists of the following parts: growing plants in specialized pots (**Fig 2A**) with interfaces enabling fitting of a cuvette covering the shoot and connection to a gas-exchange and tracer application system (see also **Fig 2C**).

**Figure 1.**
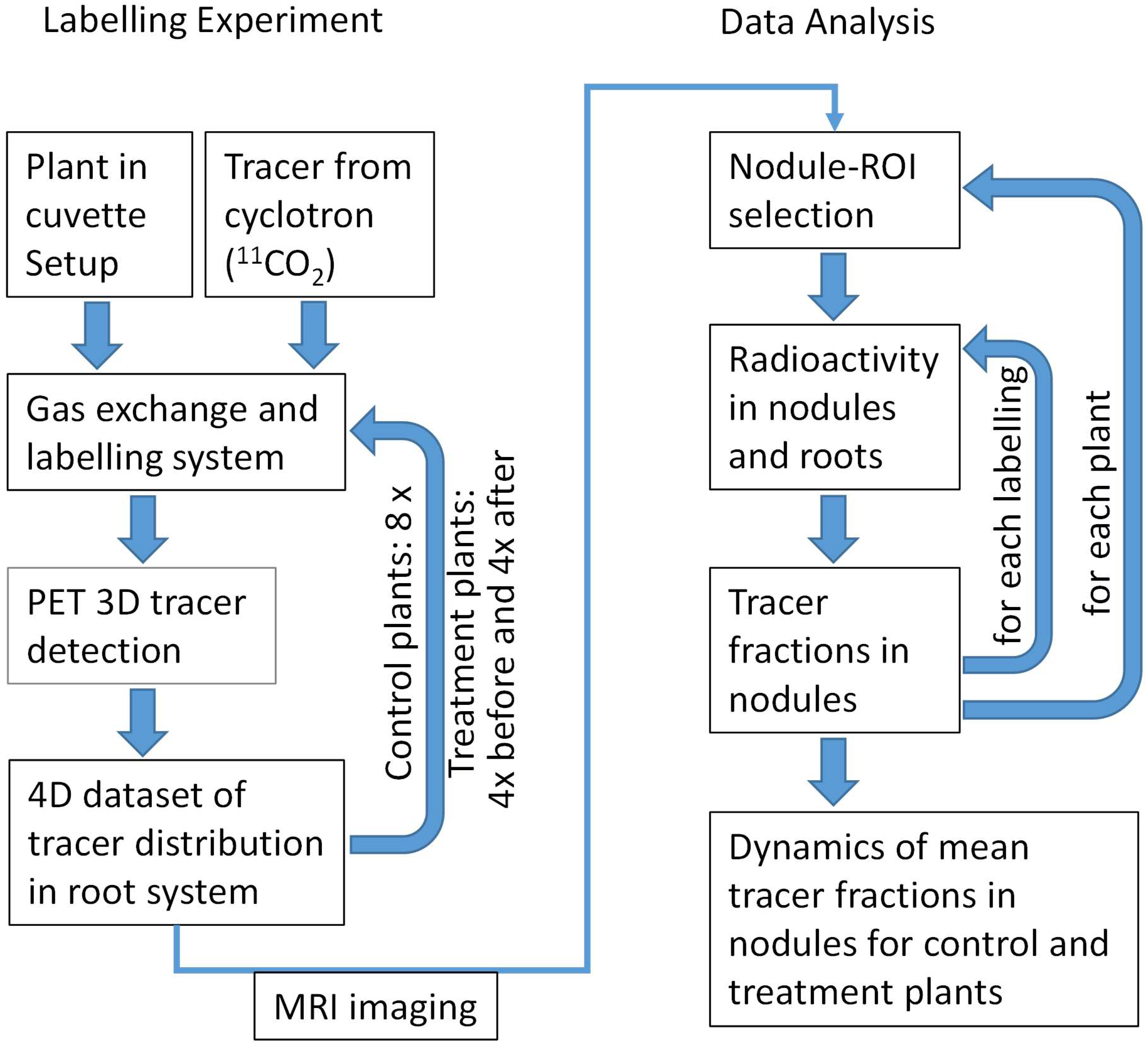
Flowchart of ^11^C-PETexperimental and data-analysis pipeline

**Figure 2:**
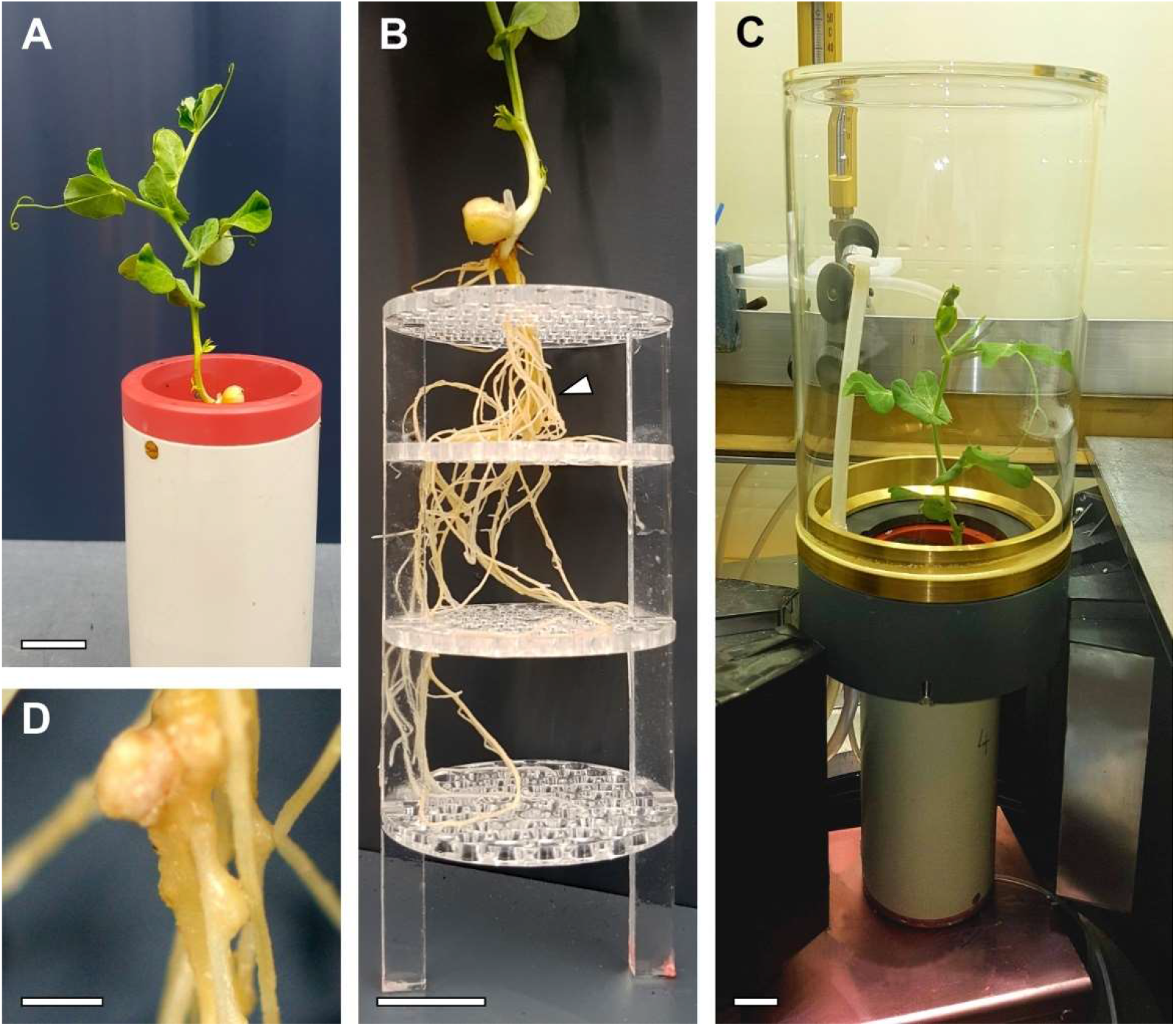
Plant cultivation and labelling set-up. **A** 16 days old Garden pea (*Pisum sativum* L. cv. Kayanne) in PVC pots with nutrient solution. **B** Transparent stabilizing support grid for reducing root displacement during plant transfer between PET and MRI measurements. Arrow indicates region shown enlarged in panel D. **C** Overview image of a pea plant with ^11^CO_2_ tracer labelling cuvette situated above the PET detectors positioned at the level of the pot. **D** Representative nodules in the root system of a pea plant. Scale bars, **A-C** 20 mm, **D** 2 mm

The cuvette enables repeated labelling of plant shoots with a 6 minute pulse of ^11^CO_2_ each time followed by measuring the ^11^C tracer 3D distribution in the root system for at least 110 minutes using a dedicated PET-system. The basic setup for the PET system in Jahnke et al. (2009). After a number of repetitions, plant root structure was analyzed non-invasively by MRI (van Dusschoten et al., 2016).

Experiments were followed by a data analysis pipeline (**Fig. 1**) consisting of MRI-assisted nodule selection and calculating the fraction of ^11^C tracer allocated to these among the whole tracer allocated to the root system from the PET data. The MRI system used here has been described in detail by . Details on the MRI and PET measurements and data analysis and quantification are given in the next section, where we describe its application to a concrete research question.

### Applying the experimental pipeline to Quantification of dynamic ^11^C allocation in individual nodules

The root systems of the pea plants were imaged with both, MRI and PET, to obtain complementary information. MRI provided a detailed 3D image of root and nodule structure which allowed to separate several well-developed nodules within the root system (**Fig. 3**A). The identification of nodules was also confirmed by optical inspection at harvest (comparable to **Figure 2**B, D). Co-registration of MRI images with PET images (**Fig. 3**B) revealed ^11^C tracer distribution within the root system. ‘Hot spots’ of tracer confirmed that large nodules corresponded to strong carbon sinks. Other areas of high tracer signal were mainly found along the primary root and the upper lateral roots (**Fig. 3**B). Regions of interest (ROIs) were defined around single prominent nodules (**Fig. 3**A, B) or small nodule clusters on the same root. Special care was taken to ensure that as few as possible root signal was included in the ROIs.

**Figure 3.**
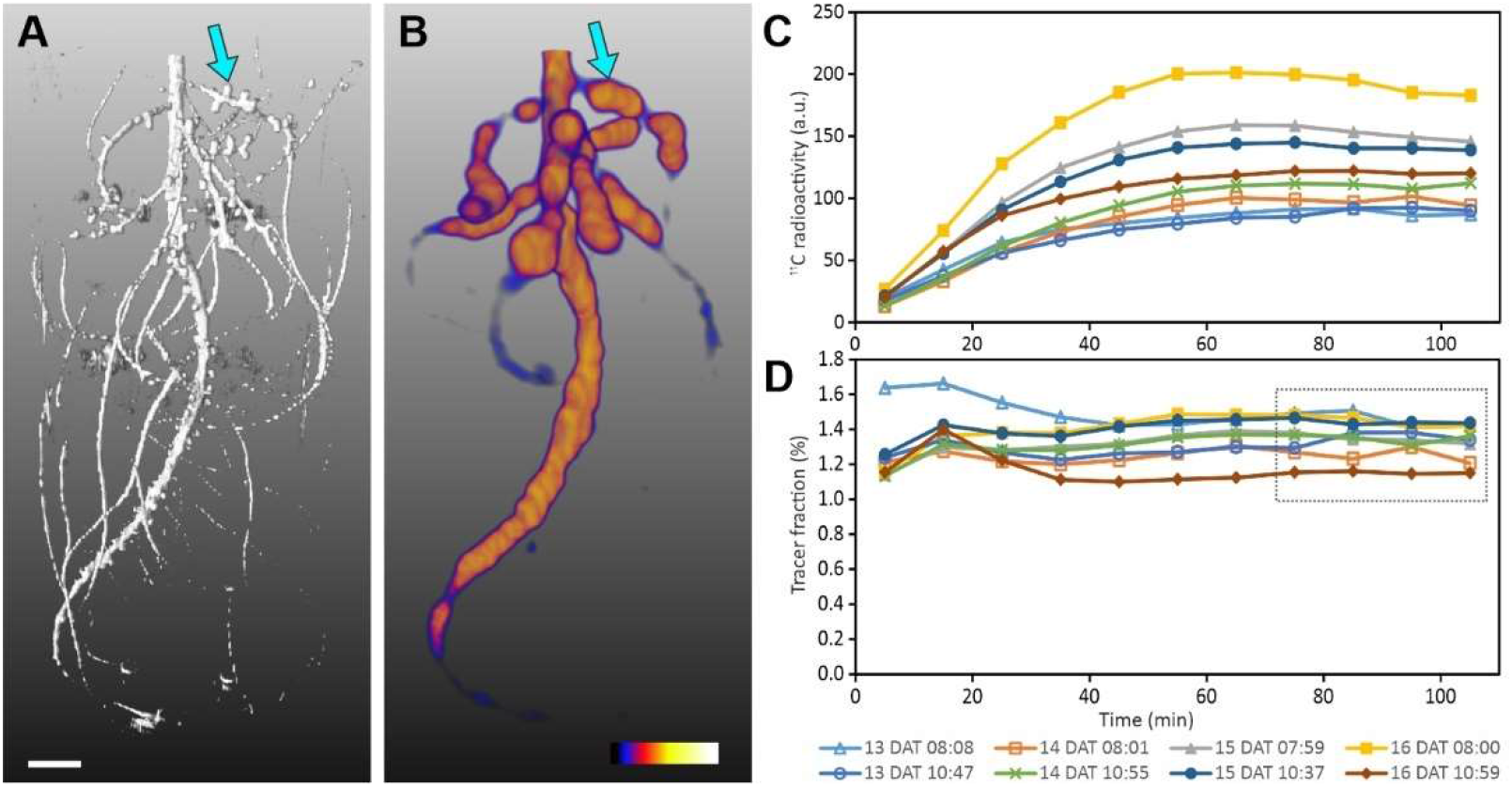
Root system of a pea plant visualized by Magnetic resonance image (MRI) and Positron Emission Tomography (PET) with corresponding time series of ^11^C tracer in one nodule ROI. **A** MRI shows the structure of a nodulated root system 16 days after transfer (DAT). The arrow indicates an individual nodule for which data are shown in C and D. Scale bar, 10 mm. B PET image of ^11^C tracer distribution within the nodulated root system 13 DAT (maximum intensity projection). Colourscale 0.0 (black) – 1.0 (white) a.u. The arrow indicates the same nodule as in A. C Time series of decay corrected ^11^C radioactivity in the region of interest (ROI) of the nodule marked by the arrow in B for all eight consecutive measurements, i.e. two measurements per day on 13, 14, 15 and 16 DAT. D Time series of radioactivity in the same ROI normalized to the total radioactivity detected in the root system. The grey box indicates the data points used to calculate mean tracer fractions for each measurement of this ROI.

Labelling experiments were performed on four consecutive days on the same plant, with two measurements per day (**Figs. 3**C, D). The time courses of the decay corrected tracer radioactivity signal in the ROIs reached a plateau around 50 min after the end of the ^11^C pulse (**Fig. 3**C), indicating that the nodules were terminal sinks for recently fixed ^11^C. The size of the signal varied considerably between the measurements (**Fig. 3**C), which may be attributed, among other reasons, to differences in transport of the ^11^C tracer inside the plant or minimal differences in labelling strength. However, our focus was on investigating dynamical responses of the tracer allocation in specific ROIs. Therefore, we normalized the ROI signals to all tracer signal in the belowground part of the plant, i.e., the integral signal visible in **Figure 3**B. The resulting fractions of activity in the ROI to the total activity in the root system are dimensionless (percentage) values which do not require decay correction and are a measure of tracer partitioning to the ROI. The tracer fractions reveal that the variability in partitioning between different measurements of the same ROI was much lower than suggested by the absolute numbers (**Figs. 3**C, D). In addition, the fractions are a better measure for the relative ^11^C allocation among the nodules. To further simplify the comparison between different measurements for several nodule ROIs at the same time, we characterized the fraction curves of **Figure 3**D by a mean calculated from four subsequent data points with least variation in all the time series (75-105 min after start of measurement). These mean fractions were used as a standard to quantify treatment effects. An example of such mean fractions for all six ROIs and eight measurements is shown in **Fig. 4**B.

**Figure 4:**
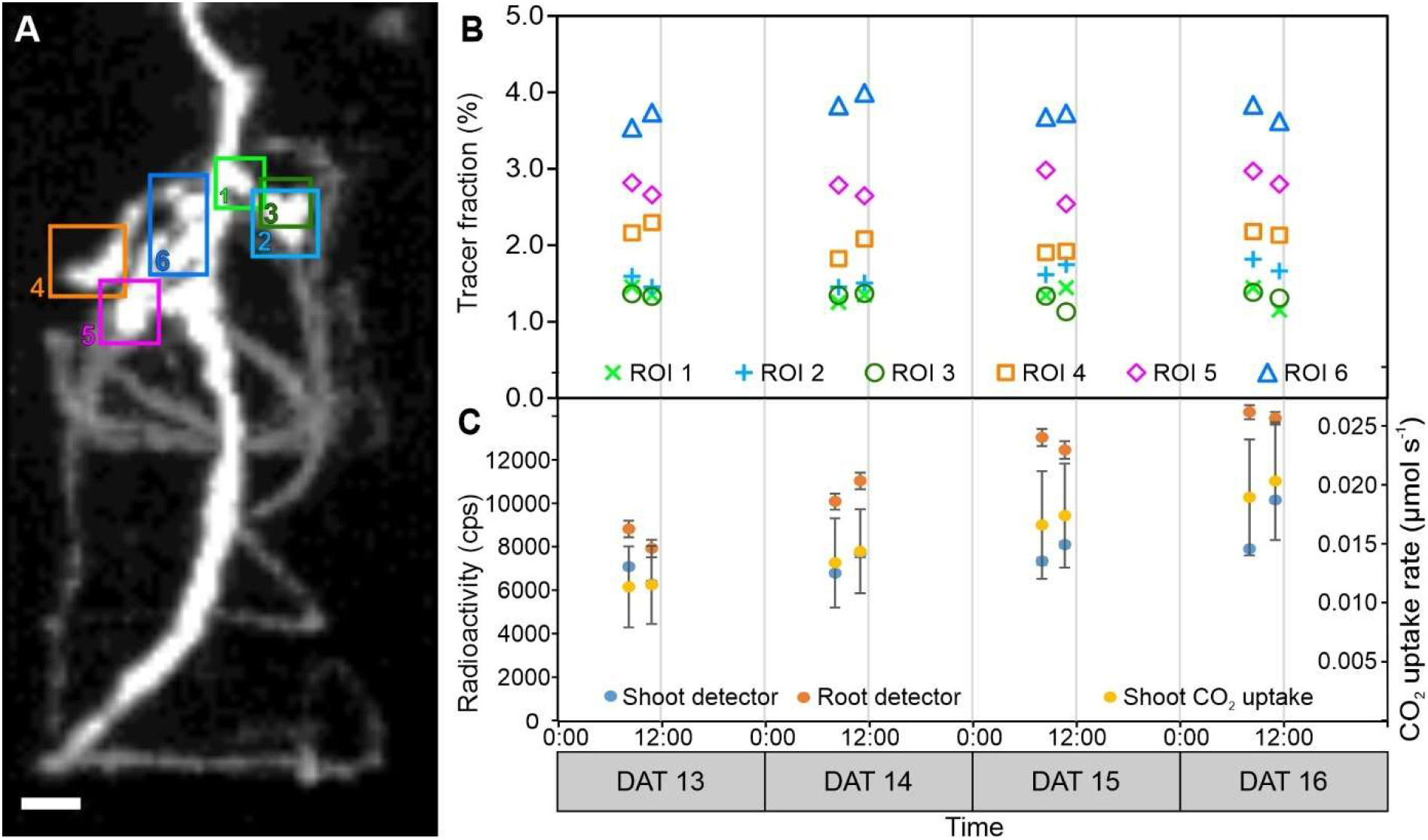
Example for carbon tracer allocation to whole root system and selected nodule ROIs of a pea (*Pisum sativum* L.) plant. A 2D-projection of carbon tracer distribution within the nodulated root system obtained from PET data, overlaid with positions of 6 ROIs enclosing selected nodules. Scale bar, 10 mm. B Mean tracer fractions in ROIs from A compared to total tracer detected in the root system, determined according to **Figure. 3**D on four days with two measurements per day. Standard deviation (SD) was lower than marker size. C Radioactivity measured by a shoot and a root scintillation detector for the same timespan the tracer fractions were calculated from (75-105 min) in counts per second (cps), decay corrected mean ±SD. Whole plant shoot CO_2_ uptake rate (mean ± SD) was determined over the full measurement period of 2h. Note that while numbers for root and shoot detector appear to be similar, the radioactivity was much higher in the shoot, where a less sensitive detector was used to prevent excessive dead time effects. DAT, days after transfer to pots.

The tracer fractions for all six nodule ROIs (**Fig. 4**A) were remarkably similar over the four days (**Fig. 4**B). There was neither a pronounced difference between the two measurements on each day nor between the different days. This is even more surprising as the plant CO_2_ uptake rate increased between 13 and 16 DAT by at least 50% (**Fig. 4**C). A similar raise in carbon uptake was indicated by an increased radioactivity monitored by the shoot detector (**Fig. 4**C). Also overall amount of radioactivity in the root increased during this time as shown by the root detector (**Fig. 4**C). Together, these data suggest that the plant did grow over the four days with increasing capability for photoassimilation. At the same time, the mean fractions of all six nodule ROIs remained similar to those on 13 DAT, indicating that the allocation pattern to the nodules was very stable over the four days and that the nodules received a constant part of the carbon allocated towards the root system.

### Effects of nitrate treatment on carbon tracer allocation towards nodules

For a more straightforward comparison between different plants and in order to compare carbon tracer allocation before and after nitrate treatment experiments, we summed up the fractions for the six nodule ROI for each plant and labelling experiment. The results are displayed in **Figure 5**, where the data points for control plant 1 correspond to **Figure 4**B. For both control plants, individual ROI fractions (**Fig. 4**B) as well as the sum of ^11^C fractions (**Fig 5**A) remained stable over the course of the experiment.

**Figure 5:**
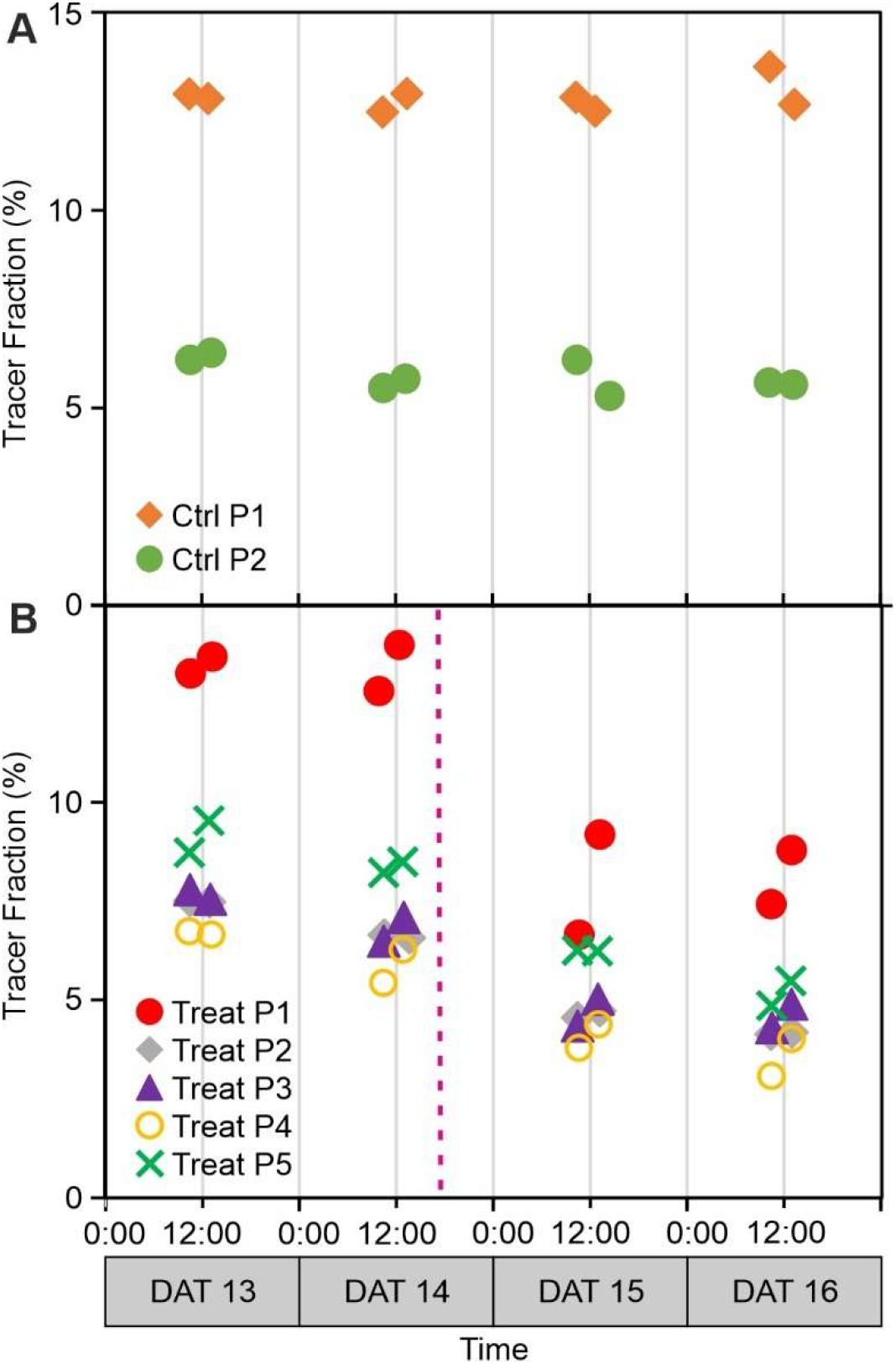
Carbon tracer allocation to nodules as fractions of the total tracer allocated to the root system detected by PET for A two control plants (Ctrl P1 and P2) and B five plants treated with nitrate (Treat P1 -5) in the root zone. Fractions for all six nodule ROIs per plant and labelling experiment were summed up to one data point. Standard deviation (SD) was lower than marker size. Individual ROI values for control Plant 1 are displayed in **Figure 4**, individual ROI values of the other plants can be found in Fig. **S2**. The starting time of the treatment is indicated by a dotted magenta line in B. DAT, days after transfer to pot.

We found an obvious difference between the tracer fractions before and after nitrate treatment in the nodules (**Fig 5**B) indicating that nitrate addition resulted in lower allocation of photoassimilates towards the nodules out of the total assimilates allocated to the root system. For statistical testing, linear slopes were calculated over all 8 measurements per plant (Table **1**). For 4 out of 5 treatment plants, the linear slopes were significantly different from zero as indicated by *p*-values < 0.05, in contrast to control plants (*p* > 0.1). The relative decrease in tracer fractions allocated to the nodules from the first day (13 DAT) to the last day of the measurements (16 DAT) amounted to 40-47% for the treatment plants (Table **1**), whereas there was no such change in the control plants. The full dataset with all treatment plants ROI can be found in the Supplemental data (Figs. **S2**, **S3** and Table **S2**).

**Table 1.** Relative change (%) of tracer fractions over the duration of the experiment.

| Plant # | linear slope<br>(% day<sup>-1</sup>) | SE<br>(% day<sup>-1</sup>) | $p$-value | Relative change (%) of tracer<br>fractions from 13 DAT to 16 DAT |
| --- | --- | --- | --- | --- |
| Control P1 | 0.07 | 0.12 | 0.314 | 2.1 |
| Control P2 | -0.20 | 0.14 | 0.135 | -10.9 |
| Treat P1 | -2.12 | 1.02 | 0.058 | -39.8 |
| Treat P2 | -1.20 | 0.50 | 0.037 | -44.6 |
| Treat P3 | -1.12 | 0.50 | 0.044 | -40.0 |
| Treat P4 | -1.08 | 0.48 | 0.046 | -47.0 |
| Treat P5 | -1.36 | 0.59 | 0.041 | -43.4 |

Linear slopes were calculated over all 8 measurements per plant. Standard errors (SE) of the estimated slopes were obtained from the least-squares regression. $p$ -values were obtained from a t-test with null hypothesis that slopes were not different from zero. DAT, days after transfer to pots

## Discussion

### Potential and challenges of using PET and ^11^C labelling to study nodules within a root system

Our study was designed to reveal feasibility of using PET with ^11^C tracer for establishing a routine in carbon allocation studies within a complex 3D structure of a root system. We found that multiple labelling of shoots with ^11^CO_2_ in combination with PET tracer detection in roots allowed for a non-invasive analysis of carbon tracer allocation towards individual nodules in the time frame of hours and days. The supplementary structural information derived from MRI measurements enabled a segmentation of individual nodule ROI to such a degree, that the dynamics of the carbon tracer signal in a small structure could be quantified as fractions of the total carbon tracer in the root system at the same time point.

To our knowledge, this is the first study that applied 3D PET to follow carbon allocation dynamics in small and localized root structures of individual plants over several days. This is a strong improvement compared to previous studies based on destructive harvests, which did not allow for temporal monitoring of individual plants e.g. Vessey et al. (1988), Voisin et al. (2003), or 1D or 2D detector approaches that were restricted to a comparison of large sections of root systems with or without nodules only (Thorpe et al., 1998, Fujikake et al., 2003, Yin et al., 2020), and also compared to earlier 3D PET work where only very limited numbers of measurements on the same individuals were perfrmed (Jahnke et al., 2009, De Schepper et al., 2012, Hubeau and Steppe, 2015).

While providing unique opportunities, application of PET and ^11^C tracer to plant studies needs to meet several challenges. The spatial resolution of PET does not always allow for unambiguous identification of small structures such as individual nodules. Co-registration with higher resolved techniques such as MRI is therefore highly valuable. However, if nodules were formed in spatial proximity or surrounded by many roots, a full separation of nodule tracer signal is not possible. Another challenge is the number of parameters that need to be controlled (i.e. kept constant) to enable comparison of repeated labelling on the same plant on several days. This includes light, temperature and humidity in the plant cuvette, but also the amount of radioactivity applied to the shoot in each labelling and gas flow through the labelling cuvette. The plant dedicated PET setup shown here met these requirements sufficiently: In the control plants the tracer fraction allocated to the nodules did not show strong differences between the different measurements. Finally, the short half-life of the ^11^C tracer (20.4 min) allows for repeated labelling experiments every few hours, but also requires a cyclotron nearby to produce the tracer. The availability of the tracer and typical length of individual measurements after labelling (2h) pose a limitation on the number of measurements per photoperiod. However, Bühler et al. (2018) provided a theoretical framework to efficiently design tracer experiments with much less PET measurement time per plant.

Also, in general the choice of substrate for the plant roots can be a challenge. The 511 keV gamma rays produced by the tracer and detected by the PET have a high penetration depth in materials such as plant substrates, but inhomogeneous absorption within the substrate may lead to problems with signal quantification. By choosing a hydroponic approach for a rapid change of nutrient conditions and facilitating root colonization by a single strain of rhizobia we avoided such problems here, as the nutrient solution is homogeneous. However, the technique can also be applied on plants potted in soil in the future which requires an additional image correction to allow quantification of the signal for inhomogeneous substrates comparable to those on clinical PET instruments.

### Application of the approach to study carbon allocation to individual nodules in response to nitrate application

The dynamics of carbon partitioning to nodules and its role in BNF are of interest, in particular with respect to environmental changes and agricultural fertilization (Schulze, 2004, Salon et al., 2011, Prudent et al., 2016). Therefore, we tested our PET approach to detect possible changes in carbon tracer allocation to nodules in response to nitrate addition. The added nitrate level was set to a level at which the BFN in nodules is expected to be reduced significantly as, applied before nodulation, it totally inhibits nodule formation (Voisin et al., 2008).

In our treatment experiments, the addition of nitrate to the root system reduced carbon tracer allocation to the monitored nodules by 40% – 47%. We determined significance of these changes by testing if linear slopes through the data points were different from zero. Even though the *p*-values were just around 0.05, the consistent difference between control plants and treated plants in the sum of nodules as well as almost all individual nodules (see Fig. **S2**) is a strong indication that the reduction of carbon allocation was a response to the nitrate treatment.

N fertilization can reduce BNF up to ~70% relative to the unfertilized control in greenhouse experiments (Santachiara et al., 2019). We suppose that BNF was reduced to a similar extent in our experiment because of the high amount of nitrate added, and notice that in spite of the reduction still a significant proportion of the carbon allocated to the nodules before treatment was reaching the nodules at least until 42h after treatment. Thus we conclude that nodules remained an important carbon sink, even at conditions where alternative and easy accessible N sources for plant uptake were available. The usage of this considerable amount of photoassimilates in the nodules, which are likely not fixing carbon any more, remains to be clarified.

Concerning the temporal dynamics of carbon, Walsh et al. (1998b) found that removing gaseous N from the surroundings of nodules for three hours did prevent nitrogen fixation, but did not affect carbon allocation to nodules. This is consistent with a decline of H_2_ evolution in Medicago starting 4h after nitrate application found by Cabeza et al. (2014). Unfortunately no data on carbon allocation are available for this experiment. Our first labelling experiment was performed 16h after start of the nitrate treatment and showed reduced allocation of carbon towards the nodules. Thus the shift may have happened in between these 4h and 16h. However due to different species and experimental conditions these data are hard to compare. This highlights that including complementary measurement of BNF in a future version of our experimental set-up might offer an opportunity to link plant carbon allocation to nodules in return to provision of N. An increased number of ^11^C pulses per day would be beneficial to gain a more detailed understanding of shorter-term responses in the first hours after N treatment and a probable recovery from a nutrient pulse as observed by Naudin et al. (2011), or whether daily rhythms interact with carbon allocation as shown for nitrogenase activity indicated by evolution of H2 (Liese et al., 2017). In the future such experiments may help to clarify whether the change in carbon allocation is a direct effect of nitrate addition that inhibited nitrogenase activity and thus reduced the carbon sink strength of nodules (Vessey et al., 1988), or if a reduced carbon allocation by the plant host served as control factor as suggested by (Schulze, 2004). In parallel, a higher frequency of measurements over a longer time period might improve understanding how a growing plant orchestrates carbon investment into rhizobial symbionts vs. into its own growth and development.

The additional monitoring of plant CO_2_ uptake as well as shoot and root radioactivity by scintillation detectors allowed us to set carbon allocation towards the nodules in the context of overall plant development. Because total activity in the root system increased over the four days of experiments in most plants (Fig. **S3**), the decrease in the tracer fractions allocated towards the nodules after nitrate treatment does not necessarily imply a reduction in the total amount of carbon available to the individual nodules. However, it indicates a clear shift in the carbon allocation pattern of the root system in disfavor of the nodules. This possibly correlates to a shift of carbon allocation towards the root tips, as several studies have found an increase in root growth stimulated by nitrate, e.g. (Fujikake et al., 2003, Ishikawa et al., 2018). Therefore, in future experiments the size of nodules could be taken into account to evaluate carbon sink strength of specific nodule tissue in comparison to root tissue, e.g. sink strength of root tips.

In conclusion, the presented ^11^C-PET-method and experimental pipeline allows for quantifying dynamics in carbon allocation in response to varying the physiological state of individual plants. The technological improvement presented here opens the door for detailed characterization of individual nodules or root types.

## Material and methods

### Plant material and pots

Seeds of garden pea (*Pisum sativum* L. cv Kayanne) were germinated on tissue paper for 6-8 days in the dark in a climate chamber with a 20 °C/16 °C (16 h/8 h) day night cycle and 60% relative humidity. For inoculation with rhizobia, a culture of the rhizobial strain p221 (courtesy of Christophe Salon, INRAe Agroecologie Dijon, France) was prepared from a stock petri dish, transferred into a Yeast-Nitrogen-Base-Medium solution and shaken (120 rpm) over night at 28 °C. 1 ml (adjusted to 0.05 optical density) of the inoculum was dripped on the roots, and the seedlings were incubated for another 24h at same conditions as for germination. Inoculated seedlings were subsequently transferred into hydroponics pots (constructed from PVC tubing, inner diameter 58 mm, height 130 mm, material thickness 2 mm; **Fig 2**) with the seeds being placed on the top grid of an acrylic support and the roots fitted through holes. In the following the age of the examined plants will be referred to as days after seedling transfer (DAT) into the pots.

Black PVC foil was used for shading the top of the pot around the seed to prevent algal growth and evaporation of the nutrient solution. The hydroponics pots were sealed at the bottom and connected to a tube (inner diameter 1 mm, B. Braun Medical AG, Sempach, Switzerland) used for both aeration and exchange of nutrient solution. The top of a pot was equipped with an interface ring that allows a direct use in the MRI robot system as described by van Dusschoten et al. (2016) (**Fig 2**A). The interface ring also served for fitting a cuvette of the tracer application and gas-exchange measurement system airtight to the pot. For keeping the roots in the same position during and between measurements, four laser-cut acrylic grids (hole diameter 6 mm) were placed at three positions in the pot: at 5 mm from the top, 19 mm from the bottom and at 25 mm distances in between (**Fig 2**B). After transfer on the grid, the plants were grown in the climate chamber under similar conditions as during germination. Lighting was provided by five 400 W HPI and five 400 W SON-T lamps (both Philips, Hamburg, Germany), which alternated every 2 hours with 5 min overlap, giving PAR intensities between 350 and 450 μmol m^−2^ s^−1^ at canopy level.

At 10 DAT the plants were transferred to the PET laboratory where the plant shoot was sealed in a glass cuvette (inner diameter 100 mm, inner height 200 mm; **Fig 2**C) attached to a gas exchange measurement and tracer application system. Lighting was provided by six 400 W HPI overhead lamps (Philips, Hamburg, Germany) at a strength of 400 μmol m^−2^ s^−1^ PAR at canopy level and a 16 h/8 h day night cycle.

Plants were grown in an N-free hydroponic nutrient solution (N-) (Table **S1**) and were thus depending on N supply by a rhizobial symbiont. To keep nutrient supply constant, the nutrient solution was exchanged on 4, 6, 10, 11, 12, 13, 14 and 15 DAT, in each case 10.5 h after light was switched on. In treatment experiments, on 14 and 15 DAT the solution was exchanged against fresh N+ solution (Table S1). The main difference between N- and N+ solutions was supplemental nitrate-N by adding 5 mM l^−1^ KNO_3_ and 5 mM l^−1^ Ca(NO_3_)_2_. To compensate for the increase in K and Ca, the amount of CaCl_2_ and K_2_SO_4_ was decreased accordingly. The solutions were prepared based on Naudin et al. (2011) but modified for a higher N-concentration treatment of 15 mM l^−1^ nitrate, which is in a range known to ensure a shut-off of BNF (Voisin et al., 2008). To avoid anoxic conditions of the hydroponic solution aeration was provided by passing a minor part of the gas influx into the shoot cuvette through the bottom of the pot.

The successful colonization with rhizobia was demonstrated by well-developed nodules e.g. (**Fig 2**D). The good vigor of shoots and roots in the absence of any N source in the nutrient solution (**Figs 2**A, D) and a strong pink color on the inside of the nodules at harvest indicated that the nodules were actively providing N to the plant host.

### Gas exchange measurement and ^11^C labelling system

The cuvette enclosing the shoot of the plant (**Fig 2**C) was connected to a gas-exchange and application system via PTFE tubing. The gas exchange and ^11^C tracer application system is based on Jahnke (2001) and can be operated in two modes: an open mode allowing gas exchange measurements, and a labelling mode enabling administration of ^11^CO_2_ to the shoot. In open mode, a gas flow (1500 ml min^−1^) was used with a CO_2_ concentration of 400 ppm at a temperature of 30 °C and 40% relative humidity. The gas was provided by mixing dry air with <1 ppm CO_2_ produced by a FT-IR-75-45 Purge Gas Generator (Parker Hannifin, Haverhill, USA) and gaseous CO_2_ from a gas bottle. Before and after passing the plant cuvette, CO_2_ concentration and humidity of the air were measured with a differential infrared gas analyser (LI 7000, LI-COR Biosciences GmbH, Bad Homburg, Germany). For labelling mode, the plant cuvette was circulated only with gas from a closed circuit in which the radioactive ^11^CO_2_ was released and radioactivity was monitored by a scintillation detector (1” NaI Scionix detector, Scionix, Bunik, The Netherlands) with a multi-channel analyser (Osprey, Mirion Technologies (Canberra) GmbH, Rüsselsheim, Germany). The whole circuit was housed in a lead shielding. During the labelling experiments, the radioactive gas with radioactivity of 450 – 470 MBq was circulated for 6 minutes through the plant cuvette. Subsequently, the system was switched back to open mode, and the air leaving the cuvette passed a soda lime absorber (Soda Lime HC Atemkalk, Medisize Deutschland GmbH, Neunkirchen, Germany), so that surplus ^11^CO_2_ not taken up by the plant was removed from the plant cuvette and safely deposited until complete decay. The PET system was started after the end of the pulse labelling to prevent excessive dead-time effects from the high levels of radioactivity in the shoot during and shortly after labelling.

### Radiotracer production and handling

Online production of ^11^CO_2_ was achieved via the ^14^N(p,a)^11^C nuclear reaction by irradiation of N in a gas target with 17 MeV protons at the baby-cyclotron BC1710 of the Institute of Neuroscience and Medicine (INM-5, Nuclear Chemistry) at Forschungszentrum Jülich. The ^11^CO_2_ was collected in specially designed trapping devices (Crouzel et al., 1987) for transfer to the plant labelling circuit.

### PET data acquisition

In this work, a custom build PET system PlanTIS (Plant Tomographic Imaging System) was used. PlanTIS was described in detail by Jahnke et al. (2009) and Beer et al. (2010) along with first imaging applications. The system is a vertical PET instrument with a cylindrical field of view of 7 cm in diameter and 11 cm in height. The resolution of the reconstructed images is 61 × 61 × 95 voxels with an isotropic voxel size of 1.15 mm. The intrinsic resolution of PET using ^11^C is physically limited to approximately 1.4 mm in biological tissues due to the high kinetic energy of the emitted positrons (Phelps, 2004).

The current setup of the PET system is shown in Fig. **S1**. The instrument was placed inside an insulated housing, providing basic climate control and lighting for the plant being imaged. PlanTIS has now been equipped with a vertical moving table which can lift the plant up and down by 8 cm allowing alternating PET measurement in the upper or lower position. This increased the vertical field of view to 19 cm after image reconstruction and allowed the measurement of the pots with 13 cm inner height. Data were acquired for 110 minutes after each 5 min ^11^CO_2_ pulse labelling and 1 minute of flushing with non-radioactive synthetic air. For a complete measurement cycle, plants remained 4 days in the PET system and were labelled two times per day. Plants were subsequently transferred to an MRI system for non-invasive root structural analysis (see MRI instrumentation and imaging), and finally harvested.

### Scintillation detectors

For gaining insights into overall plant tracer uptake, radioactivity levels in the shoot and in the roots were additionally monitored with non-imaging scintillation detectors (shoot: 2” NaI detector, type 8s8/E, Harshaw, Solon, USA; root: 3” NaI detector, type 76 SEA, Quartz & Silice, De Meern, The Netherlands) with Osprey multi-channel analysers (Mirion Technologies (Canberra) GmbH, Rüsselsheim, Germany) for quality control of the tracer application.

### Data analysis

#### PET Image reconstruction

Positron emission tomography yields three dimensional images of tracer distribution in time resulting in 4D datasets. The reconstruction of images from the positron decay events detected by PlanTIS has been described by Jahnke et al. (2009) and Beer et al. (2010). Data were reconstructed with an integration time of 5 minutes for each image in synchronization with the vertical shift of the plants. For each single measurement taking 110 minutes this resulted in 11 data points from the lower part and 11 data points from the upper part of the pot. Quadratic interpolation was used to approximate the amount of total tracer in the root system at the missing time points. Images were processed and visualized in MeVisLab (version 2.8.2, MeVis Medical Solutions AG, Bremen, Germany) as described in (Jahnke et al., 2009).

#### ROI selection and tracer profile reconstruction

Individual regions of interest (ROIs) were selected using custom written MeVisLab tools. The tools enabled a ROI selection based on a combination of 2D projections and 3D visualizations of both the root system and respective ROIs. Two different types of ROI were selected: 1) The whole root system visible in the field of view, 2) Six representative individual nodules or small nodule clusters. To ensure comparability of the ROIs in subsequent analyses, nodule ROI were chosen by three criteria: i) distinct anatomy in MRI images, ii) as few as possible roots in the ROI, and iii) a strong tracer signal, indicating strong carbon sink activity. To reduce noise and background signal in these ROIs, any signal not corresponding to a root or a nodule was masked by using a region growing module in MeVisLab. Signal strength could change between measurements by part of a nodule not being entirely in the ROI in some measurements. To avoid such effects, only nodules that showed no displacement between measurements were selected for the analysis.

#### Scintillation detectors on root and shoot

Due to the vertical shift of the plant every 5 minutes, and the resulting appearance of the belowground part of the plant in the field of view of the shoot detector (or the shoot in the root detector, respectively), detector data could only be used from the time frames when the plant was in the appropriate position to calculate average radioactivity signals for these 5 minute frames. Out of these we took averaged data from the same time frame used in the calculation of mean tracer fractions from PET measurements, i.e., 75-105 min after start of PET measurements.

#### Statistical analysis

Statistical analysis was performed in Excel 2010 (Microsoft, Redmont, USA).

### MRI instrumentation and imaging

All MRI measurements for detecting nodules and root structure were performed on a 4.7 T magnet (Magnex, Oxford, UK) with a vertical bore (310 mm inner diameter) and magnetic field gradient coils (205 mm inner diameter) providing gradients of up to 300 mT/m. A radio-frequency coil with 64 mm inner diameter (Varian, Palo Alto, CA) was used. The system has been described in detail by van Dusschoten et al. (2016). All root images were acquired using a multi-slice fast spin echo inversion recovery sequence (Haacke et al., 1999). Efficient suppression of the nutrient solution signal was achieved by reducing the T_1_ relaxation time of the nutrient solution and by adjusting the inversion time individually for each measurement to the zero crossing of the nutrient solution. For reduction of the relaxation time, the FeEDTA concentration of N-nutrient solution in the pot was increased to 1 mmol l^−1^. The following measurement parameters were used for imaging: field of view=64.4 mm, resolution 0.35×0.35 mm^2^, slice thickness=0.7 mm, TE=13 ms, TR=8.3 s, typical inversion times ~200-300 ms, bandwidth=40 kHz, turbo factor=2. For image visualization and 3D representations of the MRI datasets, the software package MeVisLab was used in combination with Matlab (Mathworks, Natick,USA) and the open source Matlab toolbox AEDES (version r172, https://github.com/mjnissi/aedes). Manual co-registration of MRI and PET images was performed in MeVisLab.

## Supplemental Material

- **Fig. S1** Set-up of the PET system with auxiliary scintillation detectors
- **Fig. S2** Carbon tracer allocation to individual nodule ROI of all Control and Treatment plants
- **Fig. S3** Whole shoot CO_2_ uptake and radioactivity measured by a shoot and a root scintillation detector for all Control and Treatment plants
- **Table S1** Composition of nutrient solutions for control and treatment
- **Table S2** *p*-values for the individual ROI data shown in Fig. S3
- **Table S3** Tracer fractions for all single Nodule ROI and all plants (MS Excel file)

## Acknowledgements

We are indebted to the team of the INM-5 Baby cyclotron at Forschungzentrum Jülich, in particular Ingo Montag and Manfred Holzgreve, for providing the ^11^CO_2_ and samples of ^18^F for calibration purposes. We would also like to thank the radiotracer team of IBG-2, namely Marco Dautzenberg and Esther Breuer, for conducting the plant experiments and Johannes Kochs for various technical help. Finally, we thank Christophe Salon and his team, INRAe Agroecologie Dijon, for providing the seeds and the rhizobial strain culture.

## Data availability

See Supplemental material Table **S3**

